# Solving Anscombe’s Quartet using a Transfer Learning Approach

**DOI:** 10.1101/2022.10.12.511920

**Authors:** Kevin Bu, Jose Clemente

## Abstract

Analysis of high-dimensional datasets often involves usage of summary statistics, one of which is the correlation coefficient. These values are then used to inform downstream analysis, whether in feature selection or in subsequent construction of networks and heatmaps. Condensing pairwise scatterplots into these singular values however, often results in a loss of information. Originally proposed by F. J. Anscombe in his famous ‘Anscombe’s Quartet,’ this phenomenon has been canonically used to demonstrate the importance of plotting and the limitations of summary statistics such as correlation or variance [F.J. Anscombe, (1973) *American Statistician*. 27 (1), 17-21]. While numerous methods exist for the generation of visually distinct datasets that share similar summary statistics, the converse has not been extensively studied. To address this gap, we propose ICLUST (Image CLUSTering), an image classifier tool that can visually distinguish correlations with similar summary statistics in simulations and identify meaningful clusters in real data. Such a tool can potentially benefit those performing exploratory analysis or feature selection in a complementary fashion by identifying relationships between variables that traditional summary metrics cannot provide.

**Significance Statement:** Distilling large-scale, multidimensional datasets via analysis of pairwise relationships often employs a single value to describe the relationship between variables. However, as demonstrated through simulations, such summarization fails to retain the nuances of the data. Characteristics such as the type of relationship (linear versus nonlinear, etc.) and the spread of the data are commonly lost when using correlations. Here we propose a transfer learning framework, borrowing from image clustering and classification software, to visually classify graphs. We apply our method towards separation of scatterplots with similar correlation statistics but visually distinctive patterns in both simulations and real data, demonstrating its broad applicability.

## Introduction

Exploratory analysis of large multidimensional datasets often relies on summary statistics such as correlation coefficients for the construction of networks and heatmaps. However, the usage of such summary statistics results in the loss of information encoded in the scatterplots of pairwise relationships. Anscombe’s quartet has canonically been used to illustrate the importance of graphing and the limitations of summary statistics such as correlation or variance - Anscombe himself stated, “make both calculations and graphs. Both sorts of output should be studied; each will contribute to understanding” [1]. This is especially critical in biological fields as Pearson and Spearman correlation are the default analytical tools when performing exploratory analysis in the gene expression and microbiome domains respectively [2–4].

Several methods have been developed to generate these kinds of datasets, analogous to Anscombe’s Quartet. The Datasaurus is one such dataset, generated using either a genetic algorithm or a simulated annealing method [5, 6]. However, there is a lack of tools that can separate these plots once they have been generated. Even the more modern exploratory data analysis tools still collapse pairwise relationships into summary statistics such as the s-Corrplot package or the MIC, which like Spearman, only quantifies strength of relationship without specifying the nature of that association [7, 8]

Here we propose ICLUST, a tool that employs transfer learning based on the pre-trained VGG16 convolutional neural network. Although the model had been trained to distinguish images of cats and dogs, by extracting the last layer of the network (4096 features), we can use the pre-trained weights to distinguish images of plotted pairwise correlations in an automated fashion, thus seeking to find ‘visual’ similarities in a way that would be impossible manually. We apply this tool to the separation of pairwise correlations from simulations and real data with the hypothesis that ICLUST can visually distinguish correlations with similar summary statistics (with performance inversely proportional to noise) and identify clusters in real data, some of which would have been masked by using correlation coefficients alone as a clustering criterion.

## Results

We first applied ICLUST to Anscombe’s quartet, taking the original data and adding to each point a specified amount of noise according to a bivariate normal distribution. Five plots were created for each class at each level of noise; the resulting set of images was then passed through ICLUST and the PCA plots are shown in **Fig. 1a.** A v-measure score (VMS) for each level of noise was computed to quantitatively assess the quality of clustering in accordance with the true labels. VMS as a function of noise (orange) is shown in **Fig. 1b** with error bars reflecting the standard deviation over one hundred such trials. The baseline for comparison (shown in blue) is the VMS obtained using clustering based on distances of the Pearson correlation summary statistic alone. Consistent with our hypothesis, increasing the level of noise reduces the accuracy of clustering as plot classes begin to overlap upon visual examination (**Fig. 1c**).

**Fig. 1.**
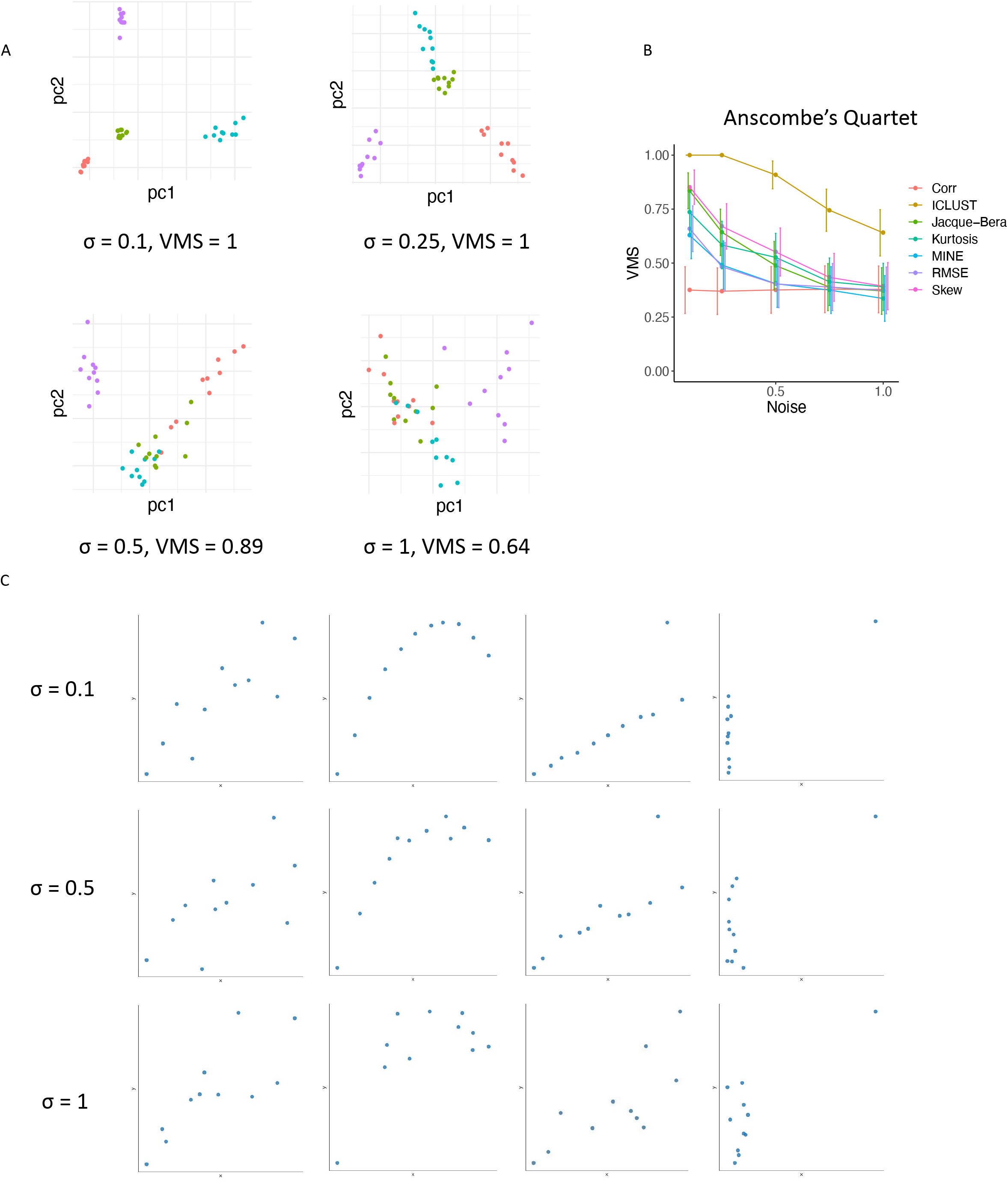
ICLUST can resolve Anscombe’s Quartet. (*A*) Principal coordinate analysis (PCA) plots (with PCA computed on the transfer learning features) at varying noise levels where points represent images of scatterplots derived from Anscombe’s quartet with the addition of noise. (*B*) V-measure score (VMS) as a function of noise level for the clustering structure in Anscombe’s Quartet (obtained via cutting the hierarchical clustering tree at k = 4). (*C*) Examples of how the plots become distorted as noise levels increase.

Given the relative efficacy of ICLUST on distinguishing clusters in canonical simulated data, we tested whether or not ICLUST could identify distinct clusters in real data. We applied ICLUST to data obtained from the WHO on a variety of health statistics for each country by computing pairwise correlations between all variables and arbitrarily choosing a window of Pearson correlation values in which to examine scatterplots. By doing so, we emulate the simulation approach described earlier, generating a dataset with similar values but potentially differing shapes and relationships. Here, we arbitrarily choose a window of correlation magnitude and select all correlations with Pearson’s *r* with a magnitude between 0.8975 to 0.9025. Hierarchical clustering based on Euclidean distance between correlation strength yields the dendrogram in **Figure 2a**, while clustering using ICLUST 4096-component feature vectors yields the structure in **Figure 2b.** Clustering assignment was determined by the best silhouette score, which corresponded to k = 2 clusters. The average image of the scatterplots in clusters 1 (red) and 2 (teal) are shown for correlation strength-based clustering and ICLUST in **Figure 2c** and **Figure 2d**, respectively. The PCA plot obtained based on Euclidean distance of the image fingerprints is shown in **Figure 2e.** Notably, variables that fall in cluster 1 tend to be normalized rates (e.g. immunization per 1000), while variables that fall in cluster 2 tend to be less uniformly distributed because of the presence of outliers. An example of this is population of a country, as countries such as India and China that are expected to be outliers skew the distribution.

**Fig. 2.**
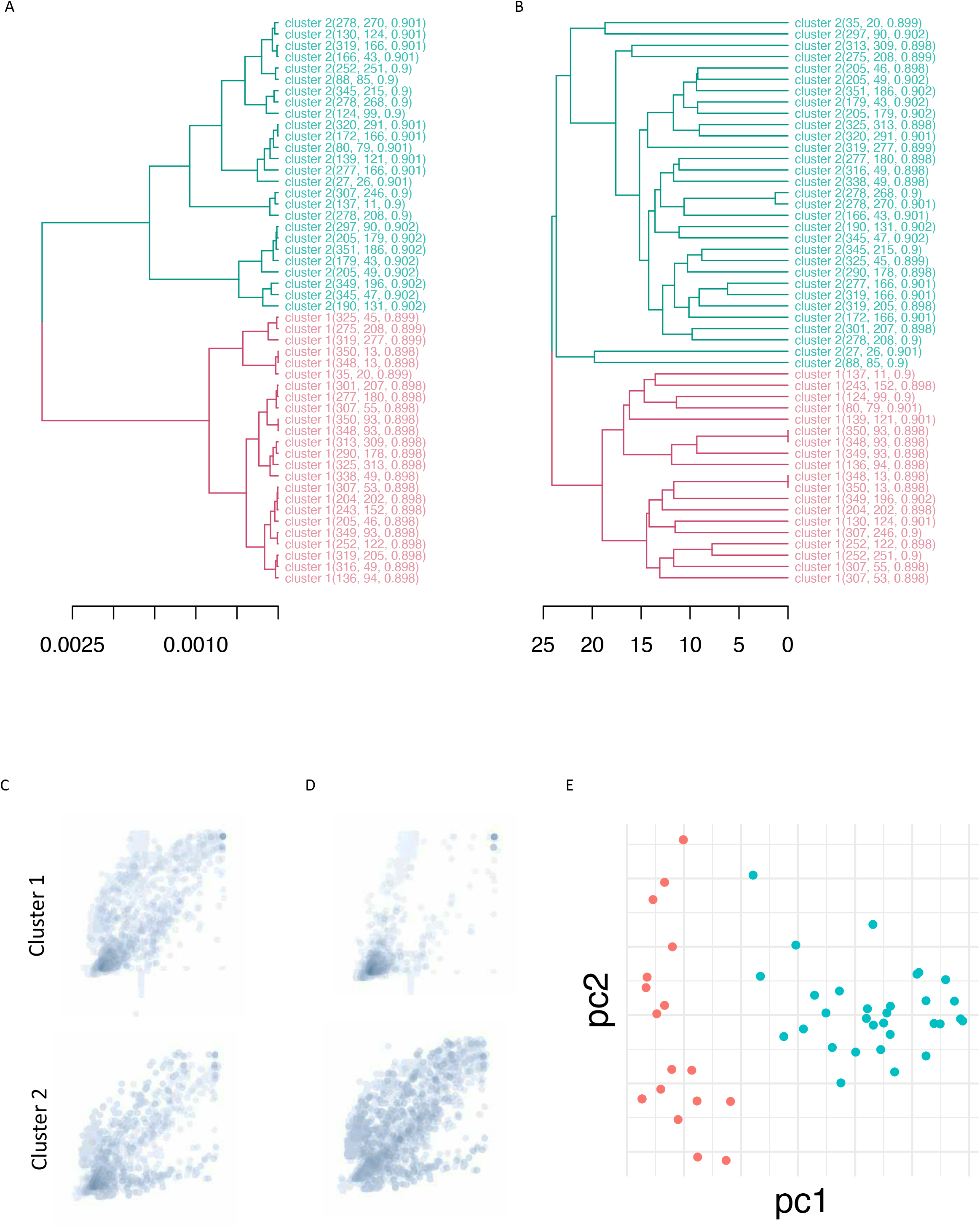
ICLUST identifies distinct clustering in WHO data. A subset of scatterplots were obtained by selecting all pairwise correlations in the WHO dataset with Pearson correlation between 0.8975 and 0.9025, with two outlier plots removed. Clustering assignment was determined by selecting the number of clusters with the highest silhouette score. (*A*) Dendrogram obtained by hierarchical clustering of scatterplots based on correlation strength alone. (*B*) Dendrogram obtained by hierarchical clustering of scatterplots based on Euclidean distance between 4096-component feature vectors of the images as processed by ICLUST. (*C*) Average image in each cluster as determined by correlation strength-based clustering, corresponding to the dendrogram in (A). (*D*) Average image in each cluster according to visual similarity clustering via ICLUST. (E). Principal Coordinate Analysis (PCA) of the scatterplots based on the 4096-component feature vector for each image with colors pertaining to the clustering obtained in (B).

We then applied ICLUST to an airline delays dataset, containing various metrics for flights (such as time spent taxiing). In this dataset, we can not only distinguish visual differences in shape (across a variety of correlation strengths, from *r* = 0 to *r* = 1) but also observe correlations that share similar correlation coefficients but distinct visual structure. When performing clustering analysis, the algorithm chooses k = 2 as the best silhouette score both when using correlation strength (**Fig. 3a**) or image fingerprints (**Fig. 3b**). The average image corresponding to these clusters for correlation strength and image fingerprints are shown in **Figure 23** and **Figure 3d** respectively. In **Figure 3c**, cluster 1 corresponds to the teal cluster in **Figure 3a** while and cluster 2 corresponds to the red cluster. In **Figure 3d**, Cluster 1 is the teal portion of the dendrogram in **Figure 3b.** The PCA plot of these clusters based on the neural network fingerprints is shown in **Figure 3e**., correlations with similar strength can appear drastically different, while correlations with different strength can appear more similar (**Fig. 3f-g**). Thus with both real examples and simulation, we demonstrate how Anscombe’s observation is indeed applicable to real world settings and that ICLUST can both separate visually distinct graphs that share summary statistics and cluster similar graphs with different correlation coefficients.

**Fig. 3.**
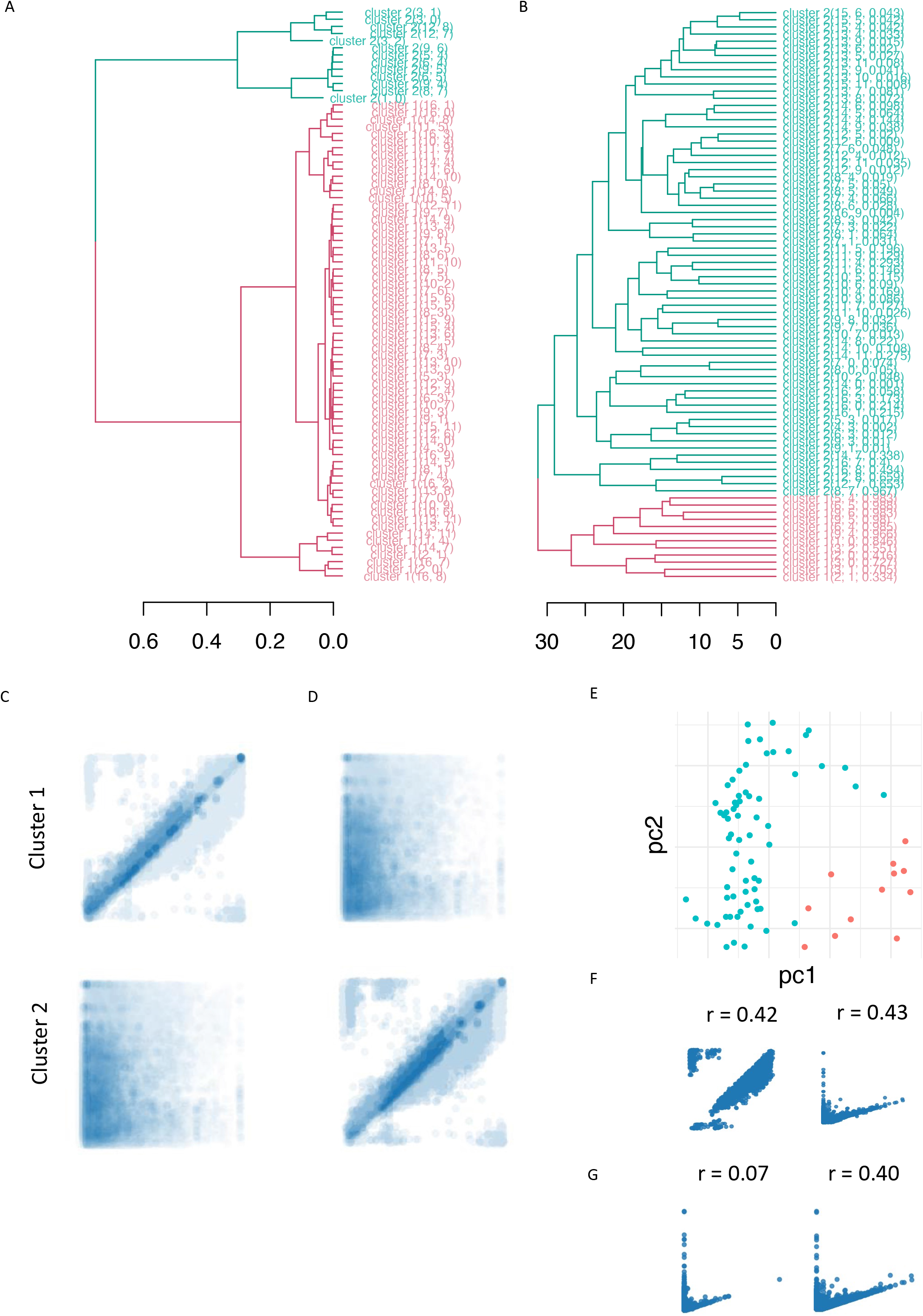
ICLUST identifies distinct clustering in airline data. All scatterplots in the dataset were plotted and clustered using (*A*) correlation strength alone and (*B*) image 4096-component feature vectors. (*C*) Average image in each cluster as determined by correlation strength-based clustering, corresponding to the dendrogram in (A). (*D*) Average image in each cluster according to visual similarity clustering via ICLUST. (E). Principal Coordinate Analysis (PCA) of the scatterplots based on the 4096-component feature vector for each image with colors pertaining to the clustering obtained in (B). (*F*) Examples of scatterplots with similar correlation size but different visual shape. (*G*) Example of correlations with similar shape but different correlation strength.

## Discussion

Given the prevalent usage of summary statistics in constructing models, networks, and other meaningful representations of data, we propose a transfer learning based image-clustering approach to the separation of scatterplots. Through simulations of Anscombe’s Quartet as well as representative real datasets (WHO, airline), we demonstrate the efficacy of ICLUST in identifying clusters of distinct patterns where summary statistics would otherwise fail to do so. Going forward, ICLUST can aid in exploratory data analysis in a complementary fashion to traditional methods, in a way consistent with Anscombe’s axiom of combining both graphs and calculations to arrive at the most accurate representation of data.

## Methods

### Plotting Simulated Data

Bivariate independent uniform displacement was added to Anscombe’s Quartet in the following manner. Let (xi, yi) be a datapoint from the dataset. We define *σ_max_* ∈ {0.1, 0.25, 0.5, 0.75, 1}; values were arbitrarily chosen to yield a representative range of noises. A new simulated dataset for each *σ_max_* was generated by computing [*x_i_* + *e*_1_*, y_i_* + *e*_2_] where *e*_1_, *e*_2_~*Unif*(0, *σ_max_*) for each dataset. Python’s Matplotlib and Seaborn libraries with were used to construct plots. Opacity of points was set to alpha = 0.1 such that overlapping points were treated differently when plotted. The default sns.lmplot function was used with palette=’set1’ and default marker size=36, shape=’o’. The origin of each plot was fixed at the center of the coordinate axes (which is hidden). The scales of the plots are allowed to vary per default plotting parameters and the method is thus scale invariant. For each set of parameters, 100 simulations were generated. Note that images shown in Fig. 1. are enlarged and include the axes for better visibility; however the clustering analysis was performed on the raw images.

### Evaluating Performance on Simulation Data

An unweighted v-measure score (VMS) was used to assess the performance of ICLUST on the labeled simulated data, as defined by:

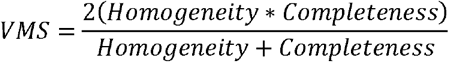

Where homogeneity is defined as:

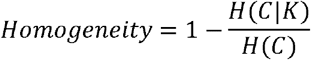

where

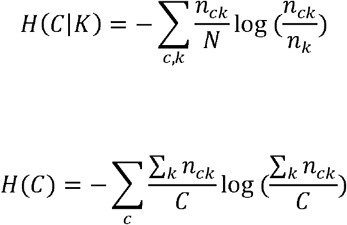

And completeness is defined as:

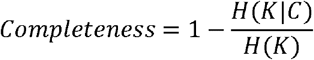

Where

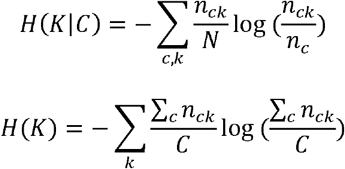

Where N is the total number of points, C is the total number of labels, and *n_c_, n_k_, n_ck_* represent the number of elements with true label C, in cluster K, and in cluster K with label C, respectively.

This is a generalization of the weighted VMS, given by:

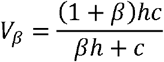

Where *β* scales the VMS by a weighting towards homogeneity; here we set *β* = 1.

### Plotting WHO and Airline Delay Data

For real-world datasets, Python’s Matplotlib and Seaborn libraries were used to construct scatterplots. Opacity of points was set to alpha = 0.1 such that overlapping points were treated differently when plotted. The default sns.lmplot function was used with palette=’set1’ and default marker size=36, shape=’o’. The upper and lower bounds for the x and y axes are dynamic and vary on a scatterplot by scatterplot basis, thus using the default parameters for determination of scaling and display. All correlations with a Pearson’s r between 0.8975 and 0.9025 in magnitude were plotted, yielding n = 51 scatterplots. The window was chosen based on a range likely to contain various shapes as described by Anscombe’s Quartet. Two plots were removed from the WHO dataset as outliers (identified via initial PCA), resulting in 49 plots. The outliers were removed to best demonstrate the two distinct clusters; visually, the outliers appeared distinct from the other plots consistent with ICLUST’s ability to distinguish visual differences between scatterplots. For the airline data, all scatterplots were plotted in the dataset (n = 80 scatterplots). WHO data and airline delay data were obtained from sources [8] and [9] respectively.

### Image Classification, Transfer Learning and Image Clustering

Image classification in ICLUST uses the VGG16 model, a convolutional neural network trained on the ImageNet dataset [10]. Briefly, input images (.PDFs,. PNGs, etc.) are scaled via the keras PIL image library which converts them into VGG16 inputs i.e. RGB (3-channel) images, each of dimensions 224 × 224 pixels. With each successive layer of the network, these pixels are converted into features using pre-trained functions. Instead of using the original output layer however, in a transfer learning setting, we adopt the penultimate layer (4096 features) as the feature map for our problem and use these as fingerprints for each image. Unsupervised clustering is performed using UPGMA after calculating the Euclidean distance between feature vectors corresponding to each image. Silhouette score is computed for each possible number of clusters, iterating from 2 through max_clust (default=10). Code for image processing and transfer learning were obtained from an open-source GitHub repository (see acknowledgements). The software was adapted from an earlier version and streamlined for use with the addition of new functionality such as concatenation of images, creation of dendrograms, generation of average images, and clustering based off of silhouette score. The average image for a given cluster is obtained by averaging the pixel intensities across the entire image for all members in the cluster. If the true class labels are given, clustering accuracy is assessed using VMS; otherwise, unsupervised clustering is performed in which the program iterates through cluster numbers (default range is 2-20) with the cluster number chosen based on the k that yields the highest silhouette score. All raw data and code used to generate analysis and figures are located at https://github.com/kbpi314/ICLUST.

## Acknowledgments

The inspiration for the software choice and architecture came from the following open-source GitHub repository by Steve Schmerler (https://github.com/elcorto/imagecluster).

